# *In vivo* imaging of the GnRH pulse generator reveals a temporal order of neuronal activation and synchronization during each pulse

**DOI:** 10.1101/2021.10.13.464231

**Authors:** Aleisha M. Moore, Lique M. Coolen, Michael N. Lehman

## Abstract

A hypothalamic pulse generator located in the arcuate nucleus controls episodic release of gonadotropin-releasing hormone (GnRH) and luteinizing hormone (LH) and is essential for reproduction. Recent evidence suggests this generator is comprised of arcuate “KNDy” cells, the abbreviation based on co-expression of kisspeptin, neurokinin B, and dynorphin. However, direct visual evidence of KNDy neuron activity at a single-cell level during a pulse is lacking. Here, we use *in vivo* calcium imaging in freely moving female mice to show that individual KNDy neurons are synchronously activated in an episodic manner, and these synchronized episodes always precede LH pulses. Furthermore, synchronization among KNDy cells occurs in a temporal order, with some subsets of KNDy cells serving as “leaders” and others as “followers” during each synchronized episode. These results reveal an unsuspected temporal organization of activation and synchronization within the GnRH pulse generator, suggesting that different subsets of KNDy neurons are activated at pulse onset than afterward during maintenance and eventual termination of each pulse. Further studies to distinguish KNDy “leader” from “follower” cells is likely to have important clinical significance, since regulation of pulsatile GnRH secretion is essential for normal reproduction and disrupted in pathological conditions such as polycystic ovary syndrome and hypothalamic amenorrhea.

## Introduction

Reproduction in mammals depends on a hypothalamic pulse generator that regulates the episodic release of gonadotropin-releasing hormone (GnRH) from the hypothalamus (*1, 2*). Regulation of the frequency and amplitude of GnRH pulses, and, in turn, that of the gonadotropins luteinizing hormone (LH) and follicle-stimulating hormone (FSH) from the anterior pituitary gland, is essential to regulate steroid hormone production and gamete development at the gonads. Although the first observations of the pulsatile nature of GnRH and LH release were made in the 1970s, until recently the precise location and cellular identity of the neural pulse generator responsible for episodic GnRH release remained a major unanswered question.

The 2003 discovery that mutations in the gene encoding the kisspeptin receptor (G protein-coupled protein 54 (GPR54)) result in hypogonadotropic hypogonadism delivered compelling evidence that kisspeptin-positive cells in the brain are required for maintaining GnRH release (*3, 4*). Subsequent studies testing the role of kisspeptin in animal models confirmed activation of GnRH neurons via GPR54 to potently stimulate GnRH and LH release (*5, 6*). Multi-labeling experiments in sheep later revealed that kisspeptin cells in the arcuate nucleus of the hypothalamus (ARC) also co-expressed two other important mediators of GnRH release; the tachykinin neurokinin B (NKB) and the endogenous opioid peptide (EOP) dynorphin (*7*); as an abbreviation, these cells were termed KNDy (Kisspeptin/Neurokinin B/Dynorphin) neurons. Colocalization of the three KNDy peptides was subsequently demonstrated in the mouse, rat, cow, goat and non-human primate (*8–12*). Anatomical characterization of KNDy neurons revealed reciprocal connections and the expression of postsynaptic receptors for NKB and dynorphin, indicating KNDy cells form an interconnected population potentially capable of synchronization (*9, 13–16*). These characteristics provided the basis for the “KNDy hypothesis” of GnRH pulse generation, in which NKB acts as the signal responsible for pulse onset by triggering activation of reciprocally connected KNDy neurons and driving the kisspeptin-mediated secretion of GnRH (*13, 17–20*). In support of this, bilateral infusions of NKB or dynorphin antagonists into the mediobasal hypothalamus (MBH) enhances and suppresses LH pulsatile release, respectively, in sheep and goats (*12, 21*). Further, the conditional inhibition and brief activation of kisspeptin-expressing cells in the ARC using optogenetic tools in mice suppresses and elicits LH pulses, respectively (*22, 23*). Finally, *in vivo* measurement of KNDy neuron population activity using GCaMP6 fiber photometry in awake and freely moving mice revealed transient increases in intracellular calcium by KNDy neurons before an LH pulse, indicative of episodic activity within the KNDy neuron population that drives pulsatile LH release (*22*). However, as these studies either manipulate or record the activity of large proportions of the KNDy population, they do not identify individual cells which are responsible for GnRH/LH pulse generation, or whether those cells are activated homogeneously during an individual pulse. To address these questions, we conducted *in vivo* calcium imaging using miniature microscopy in order to visualize KNDy cell activity at single-cell level in freely behaving mice, and, combined this with serial blood sampling to examine the pattern of activation of KNDy cells during an LH pulse. Using this approach, we showed that synchronized activation of KNDy cells always precedes an individual LH pulse. Surprisingly, we also found that not all KNDy cells are synchronized simultaneously during a pulse; rather KNDy cells are recruited in a recurring temporal order comprised of distinct subpopulations of “leader” cells, which activate and reach peak activity at the onset of synchronized episodes, and “follower cells”, which reach peak activity during either the maintenance or termination phase of the episode. These results provide an important missing piece in the “KNDy” hypothesis of pulse generation, showing that synchronized activity of individual KNDy neurons precedes each pulse, as well as revealing an unexpected temporal complexity in the cellular organization of this neural pulse generator.

## Results

### Confirmation of GCaMP6 expression in ARC kisspeptin cells

The viral vector used in this study (AAV9-CAG-Flex-GCaMP6s) contains the genetically encoded calcium sensor MP6s (GCaMP6s) and has been previously characterized as highly specific and selective to transfection of kisspeptin cells when injected into the ARC of Kiss1-Cre mice (*22*). The viral titer was diluted in this study to elongate the experimental timeline without toxicity to cells due to long-term transfection. To confirm selectivity of the viral vector to ARC kisspeptin cells, we unilaterally injected the Cre-dependent viral vector into the ARC of Kiss1-Cre/tdTomato reporter mice (Figure 1 A). We show here that 98.4 ± 0.5% of GCaMP6s-expressing cells were colocalized with tdTomato-expressing ARC kisspeptin cells, indicating, as in previous reports, the AAV is highly specific to Cre-positive cells (Figure 1 B, n=3). Conversely, 70.5 ± 8.1% of tdTomato-expressing kisspeptin cells expressed GCaMP6s, showing over two-thirds of the KNDy population express the transgene (Figure 1 C, n=3). No GCaMP6s-positive cells were detected in wild-type mice, consistent with Cre being required for transfection (n=3).

**Figure 1.**
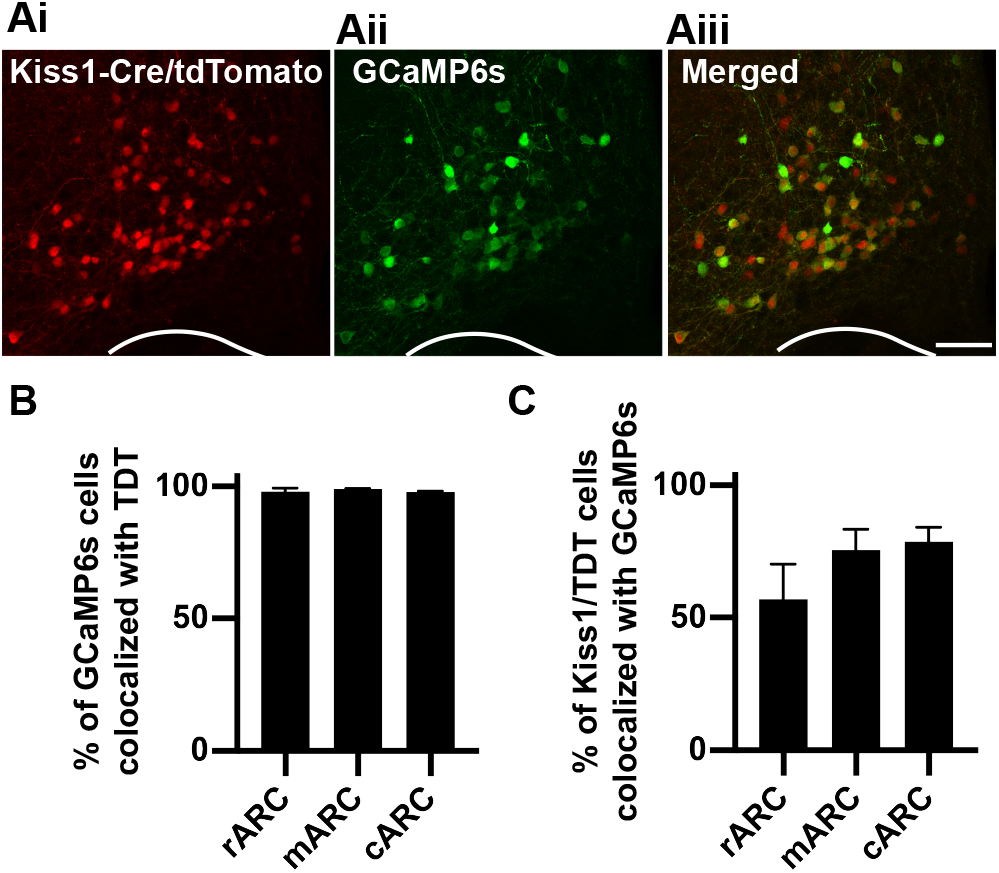
Viral delivery of the calcium indicator GCaMP6s is highly specific to arcuate kisspeptin neurons. A) Representative projected confocal images of the middle arcuate nucleus containing Kiss1-Cre cells expressing the tdTomato fluorophore (i, red), cells transfected with a Cre-dependent viral vector containing the calcium indicator GCaMP6s (ii. green) and merged images illustrating the high degree of colocalization between kisspeptin cells and GCaMP6s (iii). B-C) Histograms depicting the percentage of GCaMP6-positive cells that are colocalized with Kiss1-Cre/tdTomato throughout the rostro-caudal extent of ARC: in the rostral (r), middle (m) and caudal (c) portions (B) and the percentage of tdTomato-expressing kisspeptin cells that express GCaMP6s. Scale bar = 100μm.

### The KNDy population at a single-cell level exhibits synchronized increases in intracellular calcium

For *in vivo* calcium imaging experiments, Kiss1-Cre female mice were bilaterally ovariectomized (OVX) to remove inhibition of LH pulse generation by gonadal steroid hormone negative feedback and underwent Cre-dependent AAV transfection of ARC kisspeptin (KNDy) cells with GCaMP6s. To enable optical access to GCaMP6s-expressing KNDy cells, a 500μm diameter, 8.4mm gradient index refractive (GRIN) lens was positioned directly above the ARC and secured via an integrated baseplate. After five weeks of daily handling and habituation, unanesthetized and freely moving OVX Kiss1-GCaMP6s females underwent 60-minute sessions in which calcium fluorescent signal was visualized and recorded at 10hz via attachment of a single-photon miniaturized microscope (Inscopix Inc, nVoke) to the integrated baseplate, while tail-tip blood samples were collected every 3-4 minutes for LH pulse detection (n=5, Movie S1A, Figure 2A-C). Histological examination at the completion of recording in perfusion-fixed tissue revealed that the majority of cells recorded were located within the middle region of the ARC. Fluorescent traces were extracted and analyzed from an average of 13 ± 1.6 cells per animal using the principal component analysis (PCA)-individual component analysis (ICA) cell detection algorithm (Movie S1B-C, Figure 2D). The average reduction in baseline fluorescence during the first and final 100 seconds of baseline activity was 1.8 ± 1.4%, suggesting little to no photobleaching of cells occurred within the 60-minute recording period.

**Figure 2.**
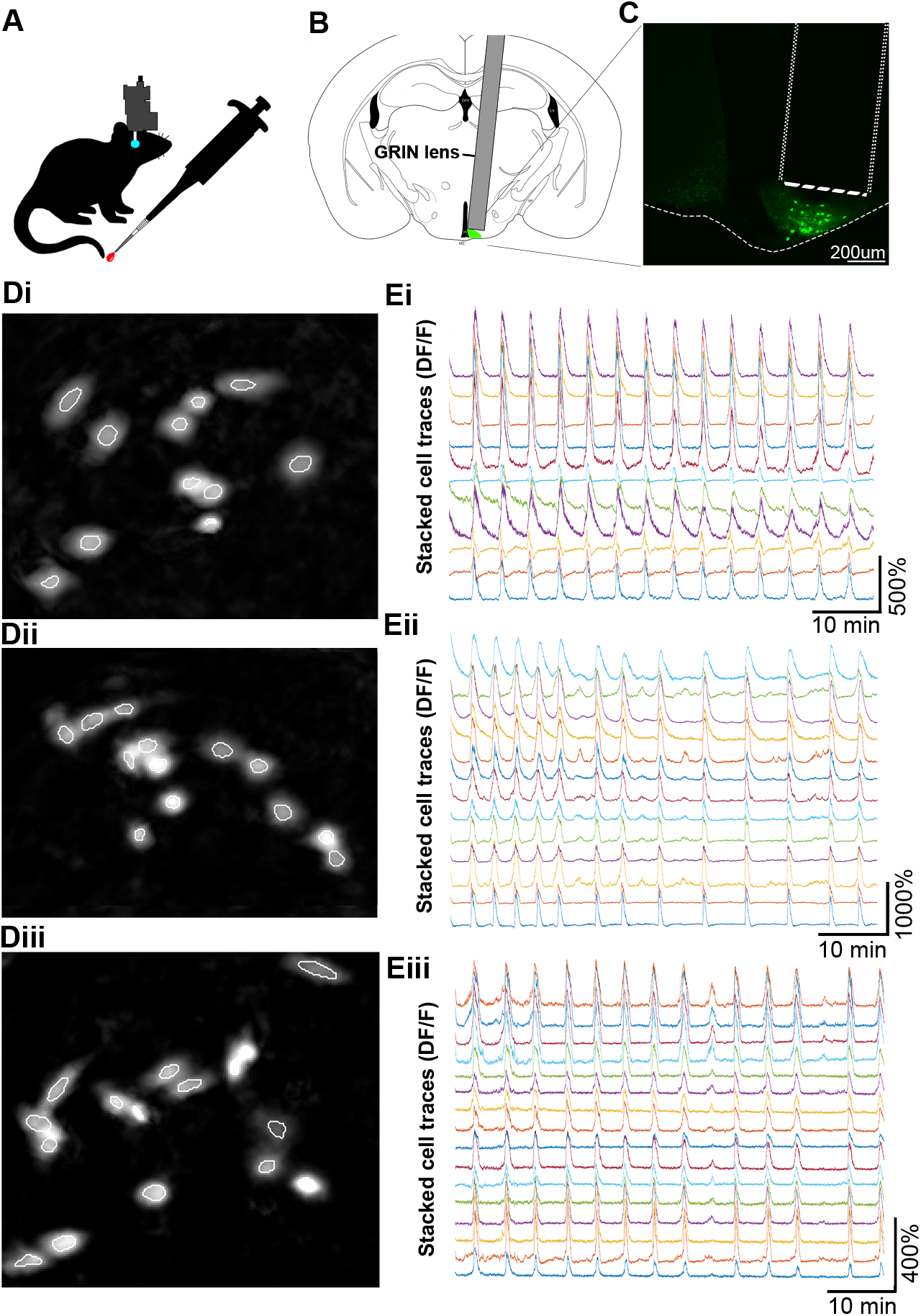
*In vivo* calcium imaging of KNDy cells in awake and freely moving mice reveals highly synchronized episodic activity at the single-cell level. A) Ovariectomized female Kiss1-Cre mice with GCaMP6s expressed by ARC kisspeptin (KNDy) cells underwent 60 minutes of *in vivo* imaging of calcium activity in the arcuate nucleus using a microendoscope (Inscopix Inc). Regular blood samples were collected from the tail-tip for luteinizing hormone pulse analysis. Schematic representation (B) and fluorescent image (C) of GRIN lens placement above GCaMP6s-expressing KNDy cells. Di-iii) Images of KNDy cells from representative animals generated following segmentation by the principal component analysis and independent component (PCA-ICA) algorithm (white outlines). Ei-iii) Representative traces of calcium activity extracted from individual KNDy cells identified using PCA-ICA in Di-iii revealing synchronized and episodic changes in fluorescence (DF/F) in awake and freely moving mice.

Visualization of KNDy neurons at the single-cell level revealed episodes of elevated calcium fluorescence across the recording period with an interval of 4.1 ± 0.7 min (Figure 2E). No observations of reduced KNDy cell activity were recorded. The average increase in fluorescent intensity relative to resting fluorescent intensity (DF/F) during episodes of activity in KNDy cells across animals was 263.98 ± 63.7%, but ranged from 117.7% to 1211.9% between individual cells. To normalize for differences in DF/F between cells, analysis was henceforth conducted on z-scored cell traces, which was generated by calculating the standard deviation of each recorded timepoint from the mean fluorescence of the cell.

Correlation analysis of single cell activity showed that although there is little to no correlation between individual cells at baseline activity, there was a strong and significant increase in correlation between cells when the average activity of the population was above baseline (Figure 3 A-B, p<0.05). In support of this high correlation during episodic activity, we detected that the majority of episodes (75.2 ± 5.7%) were the result of an increase in calcium signal by all recorded KNDy cells, representative of synchronized activation of the KNDy neuronal network and referred henceforth as synchronized episodes (SEs). In addition to SEs, episodes were also detected in which smaller percentages of the recorded KNDy cells (between 7.1-71.4%) showed episodes of elevated fluorescence, indicative of heterogenous KNDy cell activity (Figure 3C). To compare whether full versus partial recruitment of the KNDy population affected the strength and duration of network activation; the amplitude at the peak of an episode, the width of the episode at half amplitude, and the time that calcium fluorescence was above baseline was compared between SEs and episodes with only subpopulations of cells displaying activation. The average amplitude at the peak of cell activation was dramatically and significantly higher in KNDy cells during a SE compared to episodes in which only partial activation of the KNDy neuronal population was achieved (Figure 3D-E, p<0.05). Surprisingly, a positive linear relationship was not detected between the percentage of cells activated versus the amplitude at episode peak (R^2^=0.0251). Instead, except for one episode (marked # in Figure 3D), the average amplitude of the episode peak remained low unless all cells were synchronously activated. The width (s) at half amplitude, a measure of the duration of the episode, was also significantly higher during SEs compared to episodes where subpopulations were activated (p<0.05, Figure 3F). The time for cells to reach peak amplitude from baseline was not significantly different between SE and episodes with less than 100% of cells active (Figure 3G). However, the time from peak amplitude to return to baseline was significantly higher during SE’s compared to episodes with less than 100% of cells active (Figure 3H, p<0.05). Overall, the total time above baseline was significantly higher during SE’s compared to episodes with less than 100% of the recorded cells showing activity (Figure 3I, p<0.05). Together, these data indicate that recruitment of all cells during a SE result in KNDy cell activity with both higher amplitude and longer duration.

**Figure 3.**
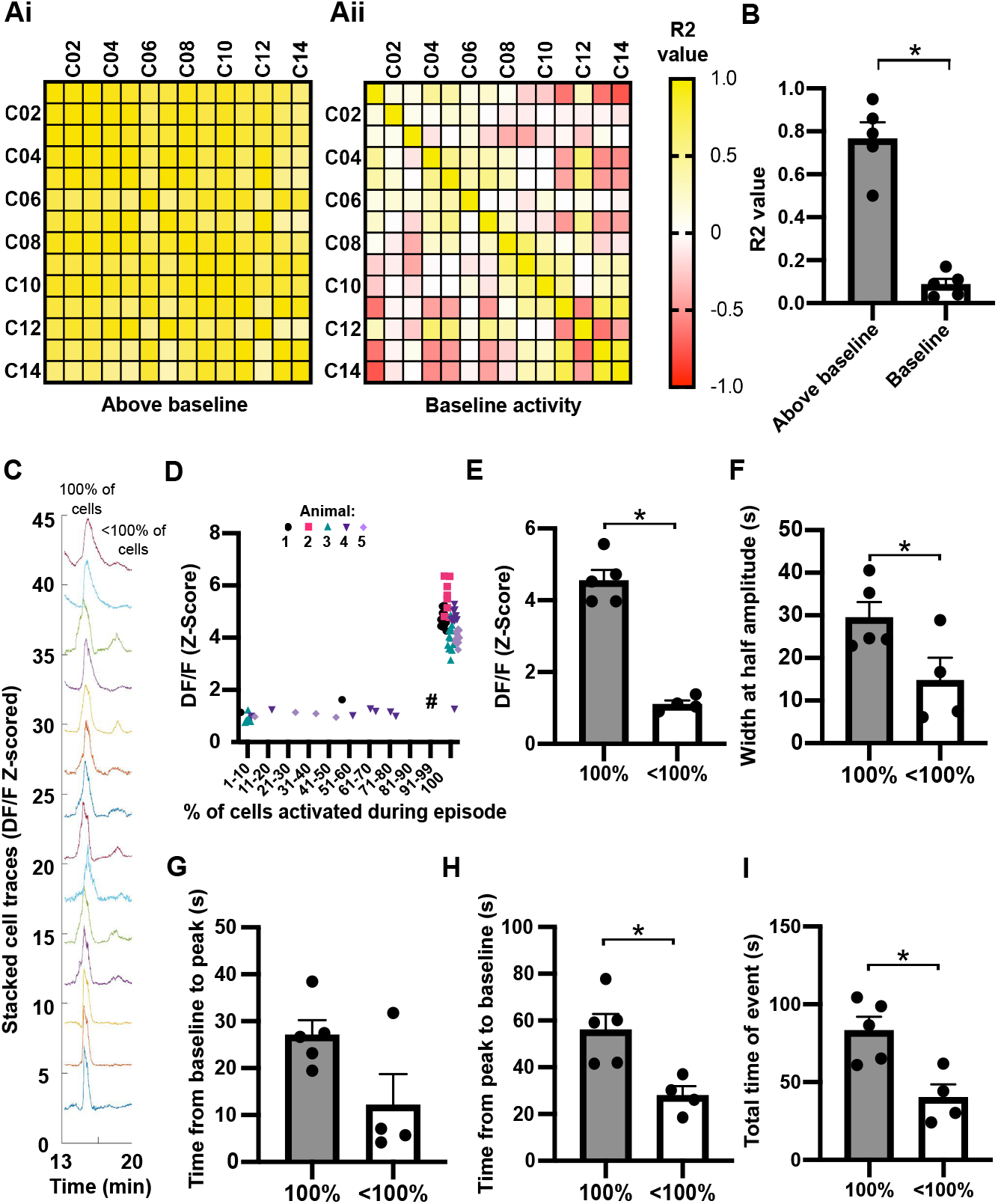
The KNDy neuron population displays either large, synchronized episodes of activation by all cells or smaller episodes of activation by subgroups of cells. A) Matrices for a representative mouse showing the correlation of activity (R^2^) between individual cells (C01-C14) when the average calcium activity for the population is above (i) or at (ii) baseline. B) The mean ± SEM R^2^ based on all animals (n=5) is significantly higher when activity is elevated above baseline. C) Representative fluorescent traces from individual cells during a synchronized episode in which 100% of cells are robustly activated, followed by lower amplitude activity by less than 100% of cells. D) Graph demonstrating the amplitude of episodes (Z-scored DF/F) remains low unless 100% of cells are activated. Each data point represents an episode in each of the animals (74 episodes from 5 animals) E-I) Graphs demonstrating the mean ± SEM (n=5) amplitude of episodes per animal (E) and the mean ± SEM width of an episode at half amplitude (F) is significantly higher when 100% of cells are activated compared to less than 100% of cells. The mean ± SEM time from baseline to peak is not significantly dependent on the percentage of cells activated (G), however the time from peak to baseline (H) and the total time above baseline (I) is significantly higher when 100% of cells are activated in an episode compared to less than 100%. * = p<0.05.

### KNDy neurons display a temporal order of activation and peak amplitude during a synchronized episode

The order in which cells 1) displayed fluorescent signal elevated above baseline, termed here as the time of cell activation (Figure 4 Ai, Bi) and 2) reached peak amplitude (Figure 4 Aii, Bii) was analyzed across all SEs recorded from each animal. Across the average of 11.2 ± 0.9 SEs recorded in 60-minutes, 45.9 ± 4.1% of recorded KNDy cells displayed at least one instance of activating first during an SE. Cells that were activating first did so on multiple SEs, on average, 2.2 ± 0.3 times out of the 11.2 ± 0.9 SEs (or 19.8 ± 0.9% of SEs) (Figure 4C). Likewise, we found that 36.6 ± 6.1% of recorded KNDy cells displayed at least one instance of reaching peak amplitude first. Cells that were peaking first would do so on an average of 3.1 ± 0.5 out of 11.2 ± 0.9 episodes (or 26.72 ± 3.2% of SEs) (Figure 4C). These data indicate the presence of a potential “leader cell” subpopulation, in which cells take turns to repeatedly activate or reach peak amplitude first. We next aimed to assess the order in which cells in this subgroup activate and reach peak amplitude when not leading the generation of an SE by activating or peaking first. To achieve this, we quantified the cumulative percentage of the cell population that was recruited (e.g. the percentage of KNDy cells that activated or peaked first or second; first, second or third; etc.) (Figure 4D). Up until the point that 33.3% of the total KNDy cell population had activated or reached peak amplitude, the size of the KNDy neuron population recruited was not significantly different from the size of the cell population that activated or peaked first. This indicates that when cells with a recorded instance of activating or peaking first are not leading an episode, they will still activate or peak early within an SE prior to two-thirds of the remaining population. An example of this can be seen in Figure 4 Bii where KNDy cells that peak first will, on alternate episodes, reach peak amplitude either second, third or fourth. Finally, we determined that only 24.9 ± 3.6% and 26.2 ± 5.3% of the recorded KNDy population exhibited at least one instance where they were last to activate or peak during the hourlong recording window, respectively (Figure 4C). Therefore, cells that were last to activate or peak in an SE did so 5.1 ± 0.9 and 5.4 ± 1.3 times out of 11.2 ± 0.9 episodes during the 60-minute recording. Only 6.2 ± 5.3 % and 4.5 ± 2.9 % of cells that activated or peaked first at least once during the recording period also displayed an instance of activating or peaking last. Together, these results suggest that KNDy cells are segregated into subpopulations of “leader” cells that either activate or peak first, and “follower” cells that either maintain or terminate the episode.

**Fig 4.**
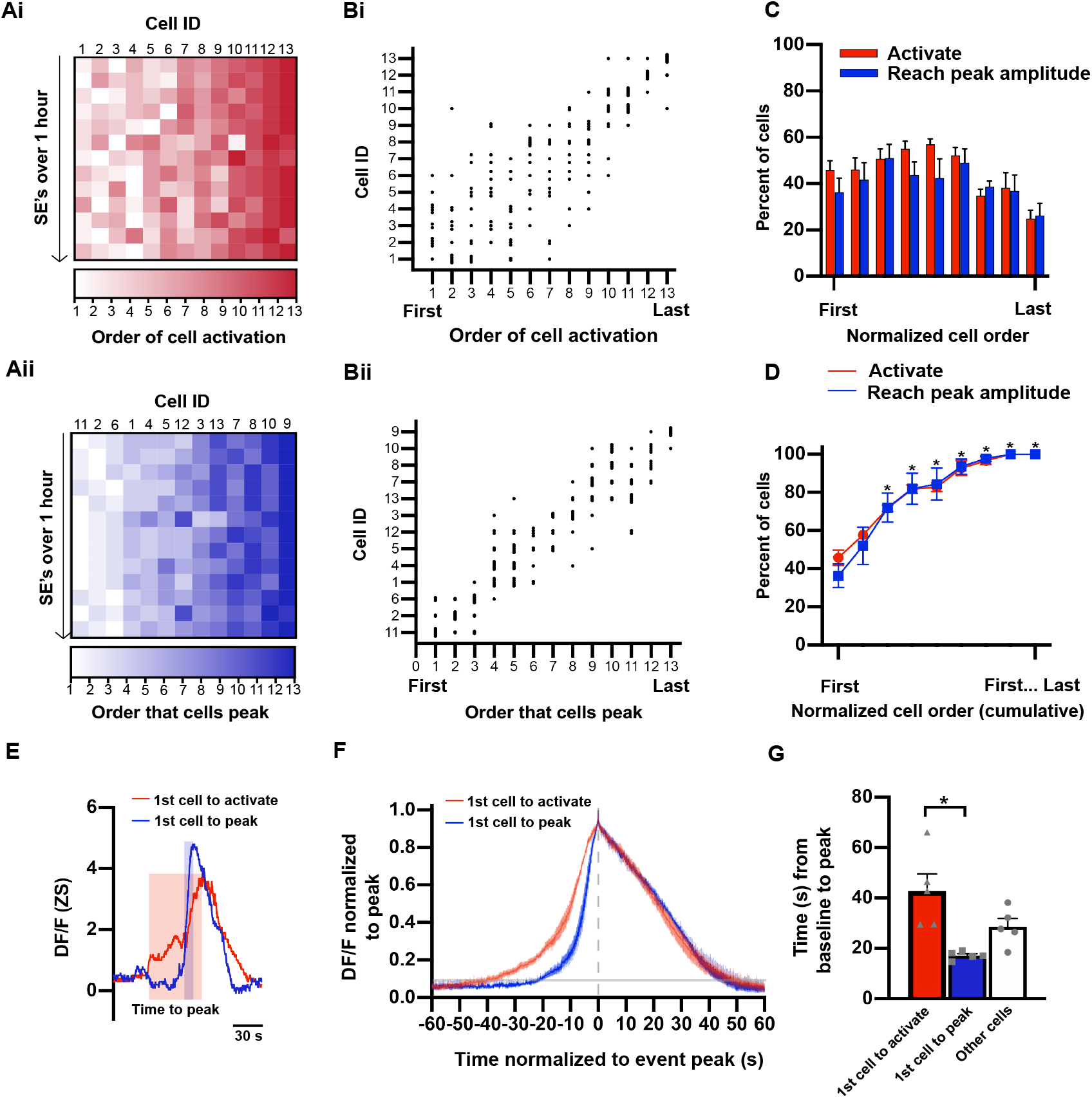
KNDy cells activate first and peak in a predictable temporal order during a synchronized episode. A) Heat-map from a representative animal depicting the order that fluorescence in individual cells (Cell ID, x-axis) elevates above baseline (i, activation) and reaches peak amplitude (ii) during synchronized episodes (y axis, 13 SE’s in 60 minutes). The cell IDs in i correspond with the cell ID’s in ii. B) Scatterplot mapping the order in which individual cells activate (i) and reach peak amplitude (ii) from the animal in A. C-D) Graphs demonstrating the mean ± SEM percentage (C) and the mean ± SEM cumulative percentage (D) of KNDy cells per animal (n=5) that activate or peak in order of first to last over multiple SEs in a 60-minute recording. * in D indicates a significant increase compared to the percent of cells that activate or peak first. E) Calcium traces from representative cells that activate first (red) and peak first (blue) during a single synchronized episode. The shaded boxes depict the time from baseline to peak for the cell that activates first (red box) and the cell that peak first (blue box). F) The mean ± SEM calcium fluorescence (normalized to peak) per animal (n=5) of cells that activate first versus cells that peak first, graphed with time 0 as the peak of fluorescent traces (grey dotted line). The shaded line on the y axis depicts the average point at which cell activity is above baseline. G) Graph illustrating the mean ± SEM time from baseline to peak amplitude per animal (n=5) for cells that activate first in an SE is significantly longer compared to cells that reach peak amplitude first. * = p<0.05.

### Synchronized episodes are initiated by a slowly activating population of leader cells

Although the percentage of KNDy cells capable of activating first was not significantly different to the population of cells that peak first (Figure 4C-D, p>0.05), only 9.1 ± 2.6% of SEs exhibited a cell that both activated first and reached peak amplitude first. Visual inspection of cell traces noted two patterns of activity that occurred between the baseline and peak of episodes. The first pattern would exhibit a rapid increase in fluorescence from baseline to peak, followed by a slower reduction in fluorescence to baseline (Figure 4E-F, blue line). The second pattern would exhibit a slower increase in fluorescence before reaching the peak of the episode (Figure 4E-F, red line). These two patterns were not observed within distinct cell subpopulations. Instead, cells could exhibit either pattern in different SEs. In line with these observations, cells that were first to activate during an SE took significantly longer, by 6.9 ± 1.5 seconds, to reach peak amplitude compared to the cells that reached peak amplitude first (Figure 4G, p<0.05.).

### LH pulses are preceded by synchronized episodes of KNDy cell activity

Serial blood samples were collected from mice throughout the 60-minute recording periods to correlate KNDy population activity with LH pulsatile release (Figure 5A). As expected in OVX female mice, a rapid frequency of 7.4 ± 0.9 LH pulses per hour was recorded. All LH pulses were preceded by KNDy neuron population activation, with an average interval of 3.1 ± 0.3 minutes lapsing between the peak amplitude of an LH pulse and the peak of the immediately preceding fluorescent episode. Although LH pulses faithfully followed KNDy neuron activation, only 49 ± 7.8% of all detected KNDy neuron activation episodes were followed by an LH pulse. Notably, LH pulses primarily followed the SEs, as 59.6 ± 6.1% of SEs (with 100% of cells activated) were followed by an LH pulse, while only one episode in which less than 100% of cells were activated (3.3 ± 3.3% of subpopulation episodes) elicited LH pulsatile release (Figure 5B). In addition, although the average interval of time between the peak of SEs (Inter-SE-Interval, ISI) across animals was 5.3 ± 0.4 min, we compared LH output within a variety of ISIs. The majority of SE’s had an ISI of over 4 minutes, although two animals exhibited episodes in which the ISI was more rapid (Figure 5C). When analyzing LH pulsatile release after KNDy SEs at varying ISIs, we found that most SEs with an ISI of over 5.5 minutes elicited an LH pulse (Figure 5D). When the ISI dropped beneath 5.5 minutes, the incidence of LH pulse generation was significantly reduced, indicating that the detection of an LH pulse depends on the ISI (Figure 5D, Figure 5Fi-iii). Effects of ISI on LH pulse amplitude were also noted. The amplitude of LH release following SEs with an ISI of 4-5.5min was not significantly different compared to episodes with an ISI of over 5.5 minutes (Figure 5E, p>0.05). However, in the two animals that exhibited SEs with an ISI of under 4 minutes and some LH pulses were detected (Figure 5D), the amplitude of the LH pulses was markedly lower compared to pulses following SEs with a larger ISI (Figure 5E). Finally, no significant difference was detected in the amplitude of SEs that were immediately followed by an LH pulse versus the amplitude of SEs that were not followed by an LH pulse (4.31 ± 0.2 vs 4.48 ± 0.4 Z-scored DF/F, p=0.7). Together, these data demonstrate that synchronized activation of the KNDy population precedes LH pulsatile release, but this relationship is not perfectly correlated when KNDy SEs are generated at high frequency.

**Fig 5.**
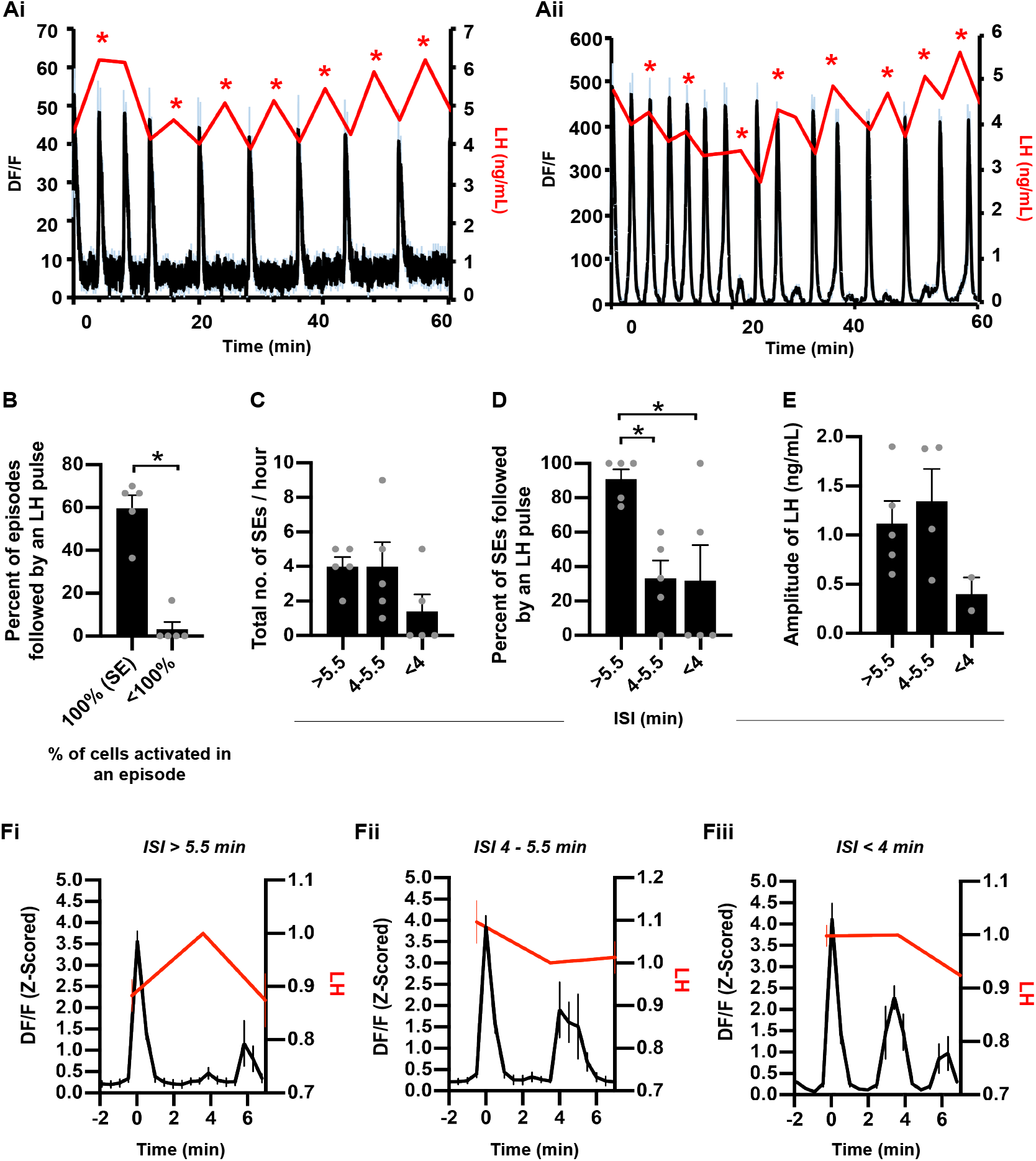
LH pulses are generated by synchronized KNDy population activity with an interval of over 5 minutes. Ai-ii) Representative examples from two animals of the mean (black line) ± SEM (blue error bars) change in fluorescence (DF/F) within the KNDy neuron population over a 60-minute recording coupled with LH pulsatile release (red line). * depicts an LH pulse. B) The percentage of episodes with KNDy cell activity followed by an LH pulse reveal episodes with less than 100% of cells recruited rarely elicit an LH pulse. C) The number of SEs over a 60-minute recording period that have an interval of over 5.5 min, 4-5.5 min and under 4 min. D) Graph summarizing that synchronized episodes (SEs) with an inter-SE-interval (ISI) of over 5.5 min are followed by an LH pulse, whereas an ISI of under 5.5 minutes significantly reduces LH pulsatile release. E) Graph illustrating that the amplitude of LH release is lower when a pulse is generated following a KNDy SE with an ISI of less than 4 minutes. F) Fluorescent traces (peak at time 0) with an ISI of over 5.5 min *(i), between 4 to 5.5 min (ii) and under 4 min (iii) plotted against LH that has normalized to the sample collected after the KNDy SE peak. * p<0.05. Data in B-F are depicted as averages per animal and expressed as mean ± SEM; n=5; * = p<0.05.

## Discussion

Using *in vivo* calcium imaging at single-cell resolution in freely behaving female mice, we demonstrate here that ARC KNDy cells, proposed as the GnRH pulse generator, show synchronous activation of all recorded cells prior to pulsatile LH release. Strikingly, we identified that the order in which cells fire during a synchronized episode is temporally organized, revealing subsets of “leaders” and “followers” that may be critical for the initiation, maintenance, and termination of GnRH pulse generation. Together, these data provide critical support that KNDy cells are the hypothalamic GnRH/LH pulse generator and novel evidence for subsets within the cell population that may dictate the initiation of activity within this reciprocally connected population.

Sixty-minute recordings of GCaMP6s fluorescent signal from the KNDy population at single cell resolution revealed that activation of all recorded cells occurred in approximately 75% of episodic events. This data supports anatomical (*14, 16, 24*) and *in vitro* electrophysiological data (*25*) which suggest KNDy cells form an interconnected network capable of synchronization. It further extends these observations by demonstrating that highly correlated synchronized activity occurs amongst all KNDy cells detected during an episode and always precedes an LH pulse. In most animals, we also detected episodes in which less than 100% of the cell population displayed activity. In these instances, the mean amplitude of the calcium signal across cells was significantly lower and not followed by LH release. Although this may indicate that complete synchronization of the KNDy population is necessary to drive LH secretion, this is not necessarily supported by previous reports. For instance, bilateral lesioning of KNDy cells can disrupt LH release (*26–28*), but the unilateral suppression of KNDy neurons using inhibitory optogenetics is insufficient to suppress LH pulsatile release (*22*). Similarly, the knockdown of approximately 30% of the KNDy cell population does not have an appreciable effect on LH pulse generation besides a slight reduction in LH pulse frequency (*29*). Combined with our results, this suggests that, *in situ,* all KNDy cells may participate in pulse generation, but not all cells may necessarily be required. Potentially, the larger amplitude recorded during endogenous synchronized episodes may be a critical factor for LH release. Currently, the role of smaller amplitude episodic events from subsets of the KNDy population is unknown. It is possible that these episodes represent failed attempts to release the necessary signal, such as NKB, believed to be a start signal for each pulse (*9, 12, 21, 30*), that drives a full synchronized episode and LH release.

Another consideration is that our data was collected from cells primarily in the middle division of the ARC. It is possible that KNDy cells in more rostral or more caudal divisions may not show the same synchronized activity before an LH pulse. For instance, previous attempts to suppress LH pulsatile release using inhibitory optogenetics was successful in the middle and caudal ARC, but not the rostral portion of the nucleus (*22*). Further, projections from the rostral half of the arcuate differ to projections from more caudal neurons (*31*). These data suggest that although we recorded from cells displaying synchronized activity consistent with pulse generation, there may be anatomically and functionally distinct subpopulations in other ARC regions that do not demonstrate the episodic activity prior to LH release and instead are related to other functions of KNDy cells, such as their role in the modulation of metabolic responses and/or stress (*32–35*).

Intriguingly, we found KNDy cells display a temporal order of firing within each synchronized episode (SE). Analysis across multiple SEs within 60-minute recording periods from each animal revealed that a subpopulation of the recorded cells peaked first at least once. When not first, this subpopulation was still temporally restricted to reach peak amplitude early within an episode. Similarly, cells that peaked during either the mid-way or endpoint of the event were restricted to this timepoint across all SEs studied. This suggests the possibility that the KNDy population contain a subset of “leader” cells capable of driving the initiation phase of an episode and “follower” cells that only fire during the maintenance and termination phase. Surprisingly, cells which were first to exhibit an increase in activity above baseline (activate) at the beginning of an episode were not the first to reach peak amplitude. Analysis of these two potential subpopulations of leader cells, cells first to activate and cells first to peak, confirmed that cells first to activate exhibited a slower ‘shoulder’ period between baseline and peak amplitude, whereas cells that peaked first demonstrated a more rapid increase in calcium activity until peak amplitude was reached. Potentially, these distinct patterns of activity may indicate a model in which “activating” leader cells undergo a slow increase in activity towards a threshold that elicits the release of NKB onto neighboring KNDy cells. NKB release may then propagate the activation of reciprocally connected KNDy cells to reach peak amplitude, as illustrated in Figure 6. The initial source of excitation for activating leader cells is currently unknown. It is worth noting, the identification of leader cells within synchronized cell populations that drive episodic pituitary hormone output has remained a major unresolved issue in multiple neuroendocrine systems. For instance, oxytocin neurons of the hypothalamus display synchronized bursts of firing that lead to pulses of oxytocin release from the posterior pituitary gland (*36, 37*). However, attempts using techniques such as paired electrophysiological recordings have been unable to determine leader and follower subpopulations (*38, 39*). Results from our data and from future studies of the KNDy pulse generator may therefore provide insight on the generation of SEs for other neuroendocrine populations as well.

Using serial blood sampling from the tail-tip during imaging, we found SEs with full activation of the KNDy cell population always occurred preceding individual LH pulses. Surprisingly, and unlike reports using fiber photometry in male mice and intact female mice (*22, 40, 41*), we also detected SEs that were not followed by an LH pulse. Further analysis revealed that the vast majority of SEs with an interval between the SE peaks (Inter-SE-Interval, ISI) of over 5.5 minutes generated a detectable LH pulse, whereas SEs with a shorter ISI weakened this relationship. Prior fiber photometry recordings detected near-perfect correlation between LH pulsatile release and preceding KNDy population activity in gonadectomized males with an interval ranging between 4.5 and 20.7 minutes (*22*). However, our observation that rapid SEs were not followed by LH release may have two explanations. First, an LH pulse with an ISI below 5.5 minutes will be difficult to capture using a blood sampling interval of 3 minutes. Here, in ovariectomized female mice, SEs displayed an average ISI of 5.3 minutes and a range between 3 and 9 minutes. The surprisingly fast pace of the GnRH pulse generator in this study may result from long-term ovariectomy which, in these studies, was typically a 6–8-week period before recording. Currently, we are unable to test whether a brief pulse of LH follows SEs with an interval of under 4 minutes, as collection of mouse tail blood samples more frequently than every 3 minutes in freely moving mice is currently not technically feasible over the length of time required for multiple SEs to occur. Alternatively, high frequency GnRH pulses may desensitize the pituitary to GnRH, leading to reduced or absent LH secretion. This is supported by historical pulse generation studies in OVX rhesus monkeys which demonstrated that using exogenous delivery of GnRH to increase the physiological frequency of GnRH stimulation of the pituitary gland from one pulse an hour to two, three and five pulses an hour resulted in a gradual decline in LH output (*42*). Similar results have been observed in perifused sheep pituitary cells, in which exogenous delivery of GnRH pulses more rapidly than every 16 minutes also reduced LH release (*43*).

A number of interesting questions arise from the results reported here: what are cellular and/or molecular features that distinguish “leader” from “follower” KNDy cells? Which signals are responsible for synchronization of “leader” cells that are first to activate versus those which are first to peak? Do manipulations of either “leader” or “follower” cells result in altered profiles of GnRH release and downstream effects on pituitary gonadotropin secretion? Given the fundamental role that the GnRH pulse generator plays in control of reproduction, identification of the mechanisms responsible for temporal ordering of KNDy cells synchronization during a pulse may have important translational relevance for understanding reproductive disease as well as normal function. In the clinic, LH pulse frequency outside the physiological norm leads to significant deficits in reproductive capacity in both male and female patients, manifesting in disorders such as hypothalamic amenorrhea (*44*) and polycystic ovary syndrome (*45*). KNDy peptide antagonists have already been intensely studied as therapeutic tools in the treatment of these and other diseases (*46–48*). Further characterization of leader versus follower cells, and the signaling pathways controlling their synchronization, may provide a basis for development of new therapeutics to either trigger or inhibit KNDy pulses and novel approaches to the control of reproduction.

## Materials and Methods

### Animals

All mice were bred and housed in the Kent State University animal facility on a 12-hour light/dark cycle and given access to food and water *ad libitum* prior to calcium imaging experiments. Experimental procedures in mice were conducted from 50 days of age. All experimental protocols and procedures were approved by Kent State University Institutional Animal Care and Use Committee under protocol 484 LC 19-08 and conform to guidelines outlined by the United States National Institutes of Health for animal research. Heterozygous Kiss1-Cre mice, in which Cre-recombinase expression is driven by Kiss1 regulatory elements (*49*) (Breeding pairs kindly donated by Dr. Carol Elias, JAX mice, stock #023426), were crossed with either C57Bl/6J mice (JAX mice, stock 000664) to generate hemizygote and wildtype Kiss1-Cre mice, or, with B6.Cg-Gt-(ROSA)^26Sortm9(CAG-tdTomato)Hze/J^ floxed-stop reporter mice (JAX mice, stock #007907) to generate heterozygous Kiss1-Cre/tdTomato female mice.

### Viral Vectors

The adenoassociated virus (AAV) pAAV2/9.CAG.FLEX.GCaMP6s.WPRE.SV40 was purchased from Addgene (Addgene viral prep #100842-AAV9; http://n2t.net/addgene: 100842; RRID: Addgene_100842). Viral titers were reduced to 2.5×10^12^ with 0.1M PBS before use. Previous studies using cell-attached recordings of ARC kisspeptin neurons in acute brain slices support that kisspeptin neuron activity is faithfully reported by changes in GCaMP6s fluorescence following transfection with the current viral vector (*22*).

### Surgical procedures

#### Stereotaxic viral injection and ovariectomy

Kiss1-Cre^+/-^, Kiss1-Cre^+/-^7tdTomato^+/-^ or Kiss1-Cre^-/-^7tdTomato^+/-^ mice were anesthetized with isoflurane (2%) and placed in a stereotaxic frame (Stoelting Co. IL, USA). Using a Drill and Microinjection Robot (Neurostar, Tubingen, Germany), a small hole was drilled into the skull 1mm posterior to bregma and 0.92 mm lateral to midline. A 29-gauge cannula attached to a 2.5μL Hamilton Syringe was loaded with 500nL of diluted AAV, angled at 6 degrees towards midline and slowly lowered 5.88 mm ventral to dura into the unilateral ARC at a rate of 150um/min. The needle was left *in situ* for 10 min before the viral vector was injected at a rate of 50nL/min. Following injection, syringes were left *in situ* for a further 10 min before the needle was slowly removed at a rate of 150um/min. During the viral injection procedure, mice were bilaterally ovariectomized to generate a state of absent steroid hormone inhibition of KNDy cell activity. Mice were returned to clean home cages and pair housed until Kiss1-Cre mice were implanted with gradient reflective index (GRIN) lenses 1-3 weeks after viral injection for calcium imaging. Kiss1-Cre/tdTomato mice were transcardially perfused with 4% paraformaldehyde in 0.1M phosphate buffered saline (PBS) 3 weeks after viral injection for immunohistochemical analysis of viral specificity with Cre-expressing arcuate kisspeptin cells.

#### GRIN lens placement

For calcium imaging, Kiss1-Cre mice injected with Cre-dependent GCAMP6s AAVs were implanted with Proview Integrated Lenses that combine a 500μm diameter, 8.4mm length GRIN lens with a baseplate (Inscopix Inc., Cat No. 1050-004611, 0.5mm diameter, 8.4mm length) designed for attachment of a miniaturized, single-photon fluorescent microscope (Inscopix Inc, nVoke). First, a 600μm diameter optical fiber (Thorlabs Inc, Cat: FT600UMT) with a sharpened tip was inserted into polyimide tubing (624μm diameter, cut to 8.4mm length, Nordson Medical, Cat: 141-0159). The polyimide-optical fiber pairing was attached to the stereotaxic frame and implanted 200-300μm above the arcuate nucleus. The implant was angled at 6 degrees towards midline to reduce motion artifacts from proximity to the third ventricle. The polyimide tubing was secured to the skull using C&B metabond dental cement (Parkell, Inc. Cat 375-0407) and, once dry, the optical fiber was retracted. The integrated GRIN lens was inserted through the polyimide tubing until 200μm above the ARC and the baseplate was secured to the skull using C&B dental cement. After surgery, mice were housed individually and allowed to recover for at least 1 week before receiving daily habituation for microscope attachment and serial blood sampling. No gross behavioral abnormalities were observed in animals that received surgery.

### Habituation to microscope attachment and serial blood sampling

One week following surgery, mice were habituated daily for 3-5 weeks to serial blood sampling protocols as reported previously (*50*). Briefly, mice were trained to enter the hand of the investigator from inside the home cage and the tail was stroked to mimic the blood collection protocol. Mice would be placed back in their home cage when samples were not being collected. In addition, mice were scruffed during handling for habituation to the restraint necessary for microscope attachment and removal.

### In vivo imaging and blood sampling

Images were acquired using a single-photon epifluorescence microscope (Inscopix nVoke miniaturized microscope, Palo Alto, CA) and Inscopix Data Acquisition Software (IDAS). Images were captured at 10 frames per second (10 Hz) with 0.8-1mW of LED (light emitting diode) power and a gain of 4. Images were collected for a total of 60 minutes per mouse. To assess LH pulsatile secretion, serial 3μL blood samples were collected from the tail-tip of freely moving mice every 3 or 4 minutes over the 60-minute imaging period and diluted in 57μL of 0.1M PBS with 0.02% Tween20. Blood samples diluted in PBS-Tween20 were immediately frozen on dry ice and then stored at −20 C until measurement of LH levels using ELISA.

### Post-processing of images

All calcium imaging data were processed using IDPS software. First, data was cropped to areas containing GCaMP6s fluorescence. Second, background was subtracted from the images using a spatial band-passing Gaussian filter. Each video was corrected for motion using the mean image of the entire video as a reference and the change in fluorescence (DF) relative to the resting F (DF/F). was generated. Within DF/F videos PCA-ICA was used to extract GCaMP6s fluorescent responses associated with individual neurons (*51*). It should be noted that while the use of automated segmentation programs to extract data from individual cells avoids bias and subjectivity in cell identification for *in vivo* calcium imaging studies, it does not perfectly detect all fluorescent cells within a recording. PCA-ICA data was manually inspected for signal from non-somatic compartments, signal detected over multiple cells, or for cells with movement that is unable to be corrected or distortion due to location on the edge of the imaging plane. Individual traces were exported for analysis in Excel and MATLAB (The Mathworks Inc.). Currently, there are two main segmentation programs used for microendoscope data: PCA-ICA and constrained non-negative matrix factorization for microendoscope imaging (CNMFe). Although CNMFe reduces the contribution of fluorescent signal from cells above and below the imaging plane when compared to PCA-ICA, it also has lower precision and identifies false positive cells (*52*). In this study, we found CNMFe had poor recognition for the somatic compartment of cells (data not shown), and therefore opted to use PCA-ICA data for analysis of calcium traces. Individual traces were exported for analysis in Excel and MATLAB (The Mathworks Inc.).

### Analysis

#### Analysis of cell activation

For all imaging experiments, potential for bleaching effects over the 60-minute imaging period were checked by comparing the baseline signal in raw traces at the beginning and end of experiments. In MATLAB, data was normalized to z-scores for quantitative analysis. Baseline fluorescence was identified using the asymmetric least square (ALS) function and was set to ‘0’ for analysis of fluctuations in calcium fluorescence. Peaks in calcium fluorescence within individual cells were identified across the 60-minute imaging period using the ‘Minpeak’ and “Mindistance” function, which detects peaks once signal surpasses 0.75 standard deviations (SD) above baseline (Z-Score of 0.75 from 0) with a minimum duration of 100s between peaks. The time and amplitude of peak fluorescence and the width (s) at half amplitude was identified. Activation and termination of an episode was defined as time when fluorescence rose above and fell below 3 times the standard deviation of the baseline of the cell, respectively. Time to peak and time to return to baseline for each episode was calculated for each cell.

#### Correlation matrix construction

To assess the correlation of activity between individual cells, correlation matrix construction was conducted using PRISM software. Z-Scored traces were separated into data points at baseline activity and data points when fluorescence reached 3 times the standard deviation of the baseline above the baseline. For both data sets, Pearson correlation was used to measure the correlation coefficient between individual cells over the 60-minute imaging period. R^2^ values from each cell-to-cell correlation were averaged per animal. R^2^ values of zero indicate a random association between neurons or no correlated activity, whereas values close to 1 or −1 indicate a high degree of positive or negative correlation, respectively.

#### Order of cell activation

To determine whether KNDy neurons fire in a predictable pattern, the order that cell activity rose above baseline (cells activated) and the order that cells reached peak amplitude was determined for each episode recorded over a 60-minute period from each animal. From this data, we calculated the percentage of the recorded cell population that displayed at least one instance of activating for each order point over the 60-minute recording session. As the number of episodes and cells recorded differed between animals, we normalized the number of cells and episodes. Further, starting from the percentage of cells that displayed at least one instance of activating or peaking first over 60 minutes of recording, we calculated the cumulative percentage of the population when including cells that activated or peaked later during the episode.

#### LH ELISA and pulse detection

Detection of LH in serial blood samples was achieved using an ultra-sensitive sandwich ELISA, as previously reported (*53*). The assay sensitivity was 0.04 ng/mL and the intra- and interassay coefficient of variation was 6.8% and 7.9%, respectively. In serial blood samples from OVX female mice, an LH pulse was identified when LH rose >10% from the previous one or two samples and lowered >20% in the following one or two samples, as previously described (*22*). The percentage of LH pulses that were preceded by KNDy neuron activation and, conversely, the percentage of KNDy population episodes followed by an LH pulse were calculated. Additionally, the time from peak of LH to the peak of the preceding episode was measured. Finally, we measured the time between peaks of adjacent synchronized episodes (Inter-SE-interval) and determined whether LH pulsatile output occurred within the interval. From this, we calculated the percentage of SE’s that were followed by an LH pulse as determined by an ISI of over 5.5 minutes, between 4 and 5.5 minutes and under 4 minutes.

#### Tissue collection and analysis of GCaMP6s specificity and expression

Following the completion of in vivo imaging experiments, mice were transcardially perfused with ice-cold 4% PFA in 0.1M PBS (phosphate buffered saline). The brain was removed from the skull and the integrated GRIN lens with cannula was carefully removed. The brain was post-fixed in 4% PFA for 1 hour at room temperature before switching to a 20% sucrose in 0.1M PBS solution until sunk. Brains were then cut into 3 parallel series of coronal sections at 30μm thickness using a freezing microtome. Tissue sections underwent 10 min washes in 0.1M PB for 3 hours and were mounted onto superfrost charged slides, air-dried and coverslipped using an aqueous mounting medium in order to image endogenous fluorescence (Gelvotol; (*54*) containing the anti-fade agent 1,4-diazabicylo(2,2)octane (Sigma-Aldrich; 50 mg/mL).

To assess the specificity of the viral vector with kisspeptin neurons, the colocalization of GCaMP6s with tdTomato cells in Kiss1-Cre/tdTomato mice was calculated. Two sections containing the rostral, middle and caudal arcuate nucleus were imaged per animal (n=3) using an Olympus FV3000 confocal microscope. A 20x objective was used to enable imaging of the entire ARC area. Optical sections with a 1.25μm step size were acquired using 488nm and 550nm channels. The number of GCaMP6s-positive cells, tdTomato cells, and dual labelled cells were quantified and the percentage of tdTomato-positive cells colocalized with GCaMP6s and the percentage of GCaMP6s-positive cells colocalized with tdTomato was calculated.

To assess viral GCaMP expression and GRINS lens placement in Kiss1-Cre mice that underwent *in vivo* Ca^2+^ imaging, tissue sections were imaged using epifluorescent microscopy (DM500B, Lecia Microsystems) and a digital camera (Microfire A/R; Optronics) paired with MicroBrightField Neurolucida Software (Williston, Vermont USA) and a 20x objective. Only mice with accurate lens placement above the ARC with viral expression of GCAMP6s displayed visible calcium signal and were included in this study. Approximately 50% of animals exhibited correct lens placement, viral infection, and fluctuations in calcium signal, and these mice were used for subsequent analysis (n=5).

#### Statistics

All statistical analyses were made using Prism8 (Graphpad Software Inc.) using the methods described below. All data, unless otherwise stated, is represented as the mean ± SEM per group (n=3 animals in Figure 1, n=5 animals in Figs. 2–6) and a p value of <0.05 was accepted as statistically significant. First, statistical comparisons of R^2^ values generated using Pearson’s correlation coefficient to determine the interaction between individual cells when at baseline activity compared to above baseline were made using two-tailed unpaired Students T-Test (Figure 3). Comparison of synchronized episodes in which 100% of cells were activated versus episodes where less than 100% of cells were activated were conducted using two-tailed unpaired Students T-Test (Figure 3). To compare the size of the recorded KNDy population that activated or peaked in order of first to last, we compared data from each order point using two-way ANOVA with *post-hoc* Bonferroni tests (Figures 4 C, D). The time between cell activation and peak amplitude for leader cells that activated first, peaked first and non-leader cells was compared using one-way ANOVA with Bonferroni post-test (Figure 4 G). Finally, the occurrence of LH pulsatile release following episodes in which all cells versus some cells were active was compared using a two-tailed Students T-test (Figure 6 B), and LH pulse generation following SEs with different ISIs were compared using one-way ANOVA with Bonferroni post-test (Figure 5 C-E).

**Fig 6.**
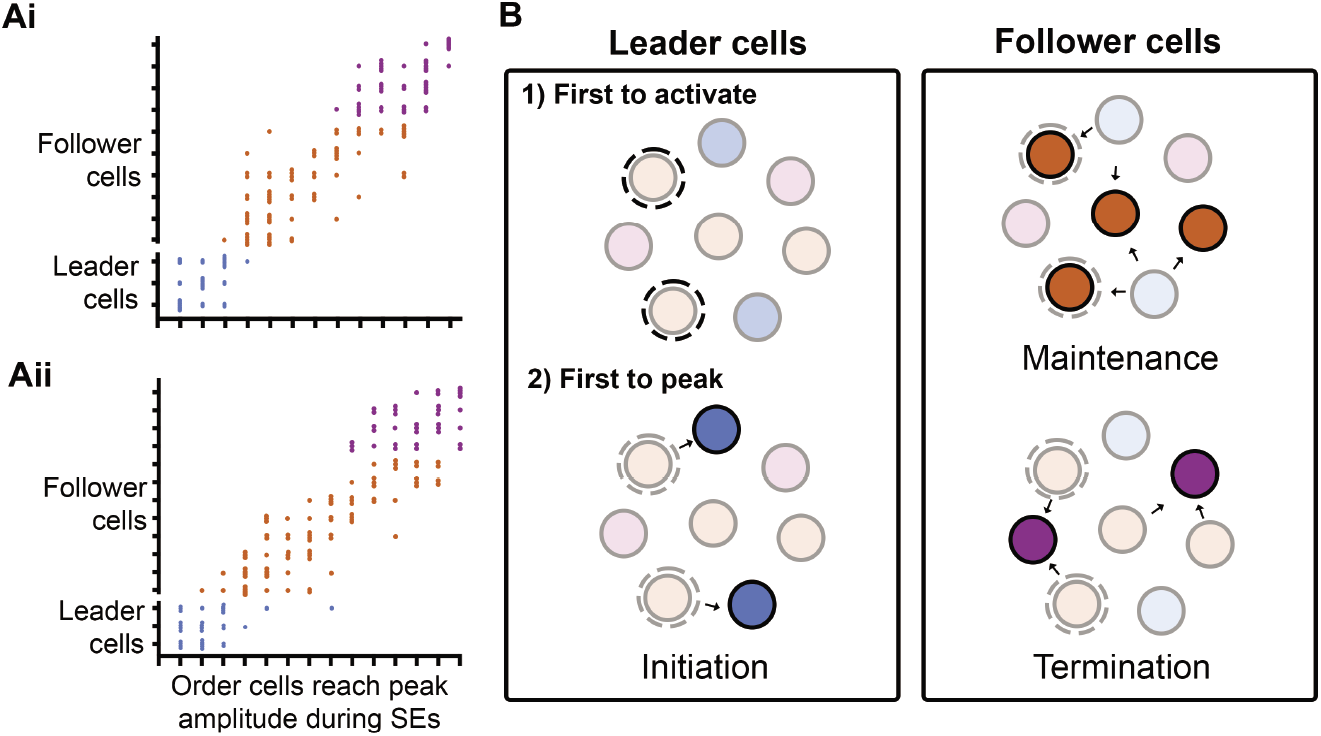
Predicted model for the temporal ordering of KNDy neuron activation during synchronized episodes. A) Scatterplots mapping the order of cell activity during synchronized episodes (SE) from 60-minute recordings from two representative animals. Each dot represents the order in which a cell reached peak amplitude during a SE. KNDy neurons have been divided into “leader” and “follower cells” depending on their order of activation across multiple SEs and color-coded into the categories shown in B. B) Schematic depicting the predicted temporal activation of KNDy neurons that drives LH pulsatile release. First, a population of leader cells first initiate an LH pulse (1) by exhibiting an increase in activity towards threshold and driving activation of a subpopulation of leader cells that peak first (2, blue cells), which activates reciprocally connected follower cells. Follower cells are further divided into cells that peak during the maintenance (orange cells) and termination (purple cells) phase of the KNDy/GnRH/LH pulse.

## Supporting information

Legend for Movie S1

## Acknowledgments

The authors would like to thank Dayanara B. Lohr for her excellent technical assistance during this research. Research reported in this publication was supported by the Eunice Kennedy Shriver National Institute of Child Health & Human Development of the National Institutes of Health under Award Numbers K99HD096120 to A.M.M and R01HD039916 to M.N.L.

## Author Contributions

A.M.M, L.M.C and M.N.L designed the research, A.M.M performed the research, A.M.M wrote the initial draft of the paper. All authors reviewed and edited the manuscript.

## References

1. Y. Nakai, T. Plant, D. Hess, E. Keogh, E. Knobil, On the sites of the negative and positive feedback actions of estradiol in the control of gonadotropin secretion in the rhesus monkey. Endocrinology 102, 1008–1014 (1978).

2. P. Belchetz, T. Plant, Y. Nakai, E. Keogh, E. Knobil, Hypophysial responses to continuous and intermittent delivery of hypopthalamic gonadotropin-releasing hormone. Science 202, 631–633 (1978).

3. N. de Roux et al., Hypogonadotropic hypogonadism due to loss of function of the KiSS1-derived peptide receptor GPR54. Proceedings of the National Academy of Sciences 100, 10972–10976 (2003).

4. S. B. Seminara et al., The GPR54 gene as a regulator of puberty. New England Journal of Medicine 349, 1614–1627 (2003).

5. M. S. Irwig et al., Kisspeptin activation of gonadotropin releasing hormone neurons and regulation of KiSS-1 mRNA in the male rat. Neuroendocrinology 80, 264–272 (2004).

6. S. Messager et al., Kisspeptin directly stimulates gonadotropin-releasing hormone release via G protein-coupled receptor 54. Proceedings of the National Academy of Sciences of the United States of America 102, 1761–1766 (2005).

7. R. L. Goodman et al., Kisspeptin neurons in the arcuate nucleus of the ewe express both dynorphin A and neurokinin B. Endocrinology 148, 5752–5760 (2007).

8. S. Ramaswamy et al., Neurokinin B stimulates GnRH release in the male monkey (Macaca mulatta) and is colocalized with kisspeptin in the arcuate nucleus. Endocrinology 151, 4494–4503 (2010).

9. V. M. Navarro et al., Regulation of gonadotropin-releasing hormone secretion by kisspeptin/dynorphin/neurokinin B neurons in the arcuate nucleus of the mouse. The Journal of neuroscience: the official journal of the Society for Neuroscience 29, 11859–11866 (2009).

10. C. True, M. Kirigiti, P. Ciofi, K. L. Grove, M. S. Smith, Characterisation of arcuate nucleus kisspeptin/neurokinin B neuronal projections and regulation during lactation in the rat. Journal of neuroendocrinology 23, 52–64 (2011).

11. A. Hassaneen et al., Immunohistochemical characterization of the arcuate kisspeptin/neurokinin B/dynorphin (KNDy) and preoptic kisspeptin neuronal populations in the hypothalamus during the estrous cycle in heifers. The Journal of reproduction and development 62, 471–477 (2016).

12. Y. Wakabayashi et al., Neurokinin B and dynorphin A in kisspeptin neurons of the arcuate nucleus participate in generation of periodic oscillation of neural activity driving pulsatile gonadotropin-releasing hormone secretion in the goat. The Journal of neuroscience: the official journal of the Society for Neuroscience 30, 3124–3132 (2010).

13. A. M. Moore, L. M. Coolen, D. T. Porter, R. L. Goodman, M. N. Lehman, KNDy Cells Revisited. Endocrinology 159, 3219–3234 (2018).

14. M. C. Burke, P. A. Letts, S. J. Krajewski, N. E. Rance, Coexpression of dynorphin and neurokinin B immunoreactivity in the rat hypothalamus: morphologic evidence of interrelated function within the arcuate nucleus. Journal of Comparative Neurology 498, 712–726 (2006).

15. P. W. Weems et al., κ-Opioid Receptor Is Colocalized in GnRH and KNDy Cells in the Female Ovine and Rat Brain. Endocrinology 157, 2367–2379 (2016).

16. C. D. Foradori, M. Amstalden, R. L. Goodman, M. N. Lehman, Colocalisation of dynorphin a and neurokinin B immunoreactivity in the arcuate nucleus and median eminence of the sheep. Journal of neuroendocrinology 18, 534–541 (2006).

17. M. N. Lehman, L. M. Coolen, R. L. Goodman, Minireview: kisspeptin/neurokinin B/dynorphin (KNDy) cells of the arcuate nucleus: a central node in the control of gonadotropin-releasing hormone secretion. Endocrinology 151, 3479–3489 (2010).

18. A. E. Herbison, The Gonadotropin-Releasing Hormone Pulse Generator. Endocrinology 159, 3723–3736 (2018).

19. H. Okamura et al., Kisspeptin and GnRH pulse generation. Advances in experimental medicine and biology 784, 297–323 (2013).

20. H. Clarke, W. S. Dhillo, C. N. Jayasena, Comprehensive Review on Kisspeptin and Its Role in Reproductive Disorders. Endocrinology and metabolism (Seoul, Korea) 30, 124–141 (2015).

21. R. L. Goodman et al., Kisspeptin, neurokinin B, and dynorphin act in the arcuate nucleus to control activity of the GnRH pulse generator in ewes. Endocrinology 154, 4259–4269 (2013).

22. J. Clarkson et al., Definition of the hypothalamic GnRH pulse generator in mice. Proceedings of the National Academy of Sciences of the United States of America 114, E10216–e10223 (2017).

23. S. Y. Han, T. McLennan, K. Czieselsky, A. E. Herbison, Selective optogenetic activation of arcuate kisspeptin neurons generates pulsatile luteinizing hormone secretion. Proceedings of the National Academy of Sciences 112, 13109–13114 (2015).

24. S. J. Krajewski, M. C. Burke, M. J. Anderson, N. T. McMullen, N. E. Rance, Forebrain projections of arcuate neurokinin B neurons demonstrated by anterograde tract-tracing and monosodium glutamate lesions in the rat. Neuroscience 166, 680–697 (2010).

25. J. Qiu et al., High-frequency stimulation-induced peptide release synchronizes arcuate kisspeptin neurons and excites GnRH neurons. eLife 5, (2016).

26. K. Beale et al., The physiological role of arcuate kisspeptin neurons in the control of reproductive function in female rats. Endocrinology 155, 1091–1098 (2013).

27. M. A. Mittelman-Smith et al., Arcuate kisspeptin/neurokinin B/dynorphin (KNDy) neurons mediate the estrogen suppression of gonadotropin secretion and body weight. Endocrinology 153, 2800–2812 (2012).

28. M. A. Mittelman-Smith, S. J. Krajewski-Hall, N. T. McMullen, N. E. Rance, Ablation of KNDy Neurons Results in Hypogonadotropic Hypogonadism and Amplifies the Steroid-Induced LH Surge in Female Rats. Endocrinology 157, 2015–2027 (2016).

29. M. Hu et al., Relative importance of the arcuate and anteroventral periventricular kisspeptin neurons in control of puberty and reproductive function in female rats. Endocrinology 156, 2619–2631 (2015).

30. T. Yamamura, Y. Wakabayashi, S. Ohkura, V. M. Navarro, H. Okamura, Effects of intravenous administration of neurokinin receptor subtype-selective agonists on gonadotropin-releasing hormone pulse generator activity and luteinizing hormone secretion in goats. The Journal of reproduction and development 61, 20–29 (2015).

31. S.-H. Yeo, A. E. Herbison, Projections of arcuate nucleus and rostral periventricular kisspeptin neurons in the adult female mouse brain. Endocrinology 152, 2387–2399 (2011).

32. J. Roa, M. Tena-Sempere, Connecting metabolism and reproduction: roles of central energy sensors and key molecular mediators. Molecular and cellular endocrinology 397, 4–14 (2014).

33. C. C. Nestor, M. J. Kelly, O. K. Ronnekleiv, Cross-talk between reproduction and energy homeostasis: central impact of estrogens, leptin and kisspeptin signaling. Hormone molecular biology and clinical investigation 17, 109–128 (2014).

34. P. Grachev et al., Neurokinin B signaling in the female rat: a novel link between stress and reproduction. Endocrinology 155, 2589–2601 (2014).

35. J. A. Yang et al., Acute Psychosocial Stress Inhibits LH Pulsatility and Kiss1 Neuronal Activation in Female Mice. Endocrinology 158, 3716–3723 (2017).

36. Y. Otsuki, K. Yamaji, M. Fujita, T. Takagi, O. Tanizawa, Serial plasma oxytocin levels during pregnancy and labor. Acta obstetricia et gynecologica Scandinavica 62, 15–18 (1983).

37. M. R. Perkinson, J. S. Kim, K. J. Iremonger, C. H. Brown, Visualising oxytocin neurone activity in vivo: The key to unlocking central regulation of parturition and lactation. Journal of neuroendocrinology, e13012 (2021).

38. V. Belin, F. Moos, Paired recordings from supraoptic and paraventricular oxytocin cells in suckled rats: recruitment and synchronization. The Journal of Physiology 377, 369–390 (1986).

39. V. Belin, F. Moos, P. Richard, Synchronization of oxytocin cells in the hypothalamic paraventricular and supraoptic nuclei in suckled rats: direct proof with paired extracellular recordings. Experimental brain research 57, 201–203 (1984).

40. H. J. McQuillan, S. Y. Han, I. Cheong, A. E. Herbison, GnRH pulse generator activity across the estrous cycle of female mice. Endocrinology 160, 1480–1491 (2019).

41. S. Y. Han, I. Cheong, T. McLennan, A. E. Herbison, Neural Determinants of Pulsatile Luteinizing Hormone Secretion in Male Mice. Endocrinology 161, (2020).

42. L. Wildt et al., Frequency and amplitude of gonadotropin-releasing hormone stimulation and gonadotropin secretion in the rhesus monkey. Endocrinology 109, 376–385 (1981).

43. R. P. McIntosh, J. E. McIntosh, Influence of the characteristics of pulses of gonadotrophin releasing hormone on the dynamics of luteinizing hormone release from perifused sheep pituitary cells. J Endocrinol 98, 411–421 (1983).

44. R. B. Perkins, J. E. Hall, K. A. Martin, Aetiology, previous menstrual function and patterns of neuro-endocrine disturbance as prognostic indicators in hypothalamic amenorrhoea. Human reproduction (Oxford, England) 16, 2198–2205 (2001).

45. C. Coyle, R. E. Campbell, Pathological pulses in PCOS. Molecular and cellular endocrinology 498, 110561 (2019).

46. J. T. George et al., Neurokinin B Receptor Antagonism in Women With Polycystic Ovary Syndrome: A Randomized, Placebo-Controlled Trial. The Journal of clinical endocrinology and metabolism 101, 4313–4321 (2016).

47. J. K. Prague et al., Neurokinin 3 receptor antagonism as a novel treatment for menopausal hot flushes: a phase 2, randomised, double-blind, placebo-controlled trial. Lancet (London, England) 389, 1809–1820 (2017).

48. C. N. Jayasena et al., Subcutaneous injection of kisspeptin-54 acutely stimulates gonadotropin secretion in women with hypothalamic amenorrhea, but chronic administration causes tachyphylaxis. The Journal of Clinical Endocrinology & Metabolism 94, 4315–4323 (2009).

49. R. M. Cravo et al., Characterization of *Kiss1* neurons using transgenic mouse models. Neuroscience 173, 37–56 (2011).

50. R. B. McCosh, M. J. Kreisman, K. M. Breen, Frequent tail-tip blood sampling in mice for the assessment of pulsatile luteinizing hormone secretion. Journal of visualized experiments: JoVE, (2018).

51. E. A. Mukamel, A. Nimmerjahn, M. J. Schnitzer, Automated analysis of cellular signals from large-scale calcium imaging data. Neuron 63, 747–760 (2009).

52. A. M. Stamatakis et al., Miniature microscopes for manipulating and recording in vivo brain activity. Microscopy (Oxford, England), dfab028 (2021).

53. F. Steyn et al., Development of a methodology for and assessment of pulsatile luteinizing hormone secretion in juvenile and adult male mice. Endocrinology 154, 4939–4945 (2013).

54. L. B. Kuiper, L. N. Beloate, B. M. Dupuy, L. M. Coolen, Drug-taking in a socio-sexual context enhances vulnerability for addiction in male rats. Neuropsychopharmacology: official publication of the American College of Neuropsychopharmacology 44, 503–513 (2019).

