## Supplementary material for "*In vivo* imaging of the GnRH pulse generator reveals a temporal order of neuronal activation and synchronization during each pulse": Legend for Movie S1

**This PDF file includes:**

Legends for Movie S1

**Other Supplementary Material for this manuscript includes the following:**

Movie S1

### SUPPLEMENTARY MATERIALS

**Supplementary Movie S1. *In vivo* visualization KNDy neuron activity in ovariectomized and freely moving female mice.** A) Recording of raw fluorescent signal from GCaMP6s-expressing kisspeptin neurons in the ARC (KNDy neurons) after motion correction. B) The change in fluorescent intensity relative to the resting fluorescent intensity (DF/F) of GCaMP6s signal in A. C) Traces of DF/F from individual cells extracted by PCA-ICA analysis in B. The recording spans one hour and is shown at 60x speed.
